# Using a decision tree to predict the number of COVID cases: a tutorial for beginners

**DOI:** 10.1101/2023.12.19.572463

**Authors:** Lucy Moctezuma, Lorena Benitez Rivera, Florentine van Nouhuijs, Faye Orcales, Allen Kim, Ross Campbell, Megumi Fuse, Pleuni S. Pennings

## Abstract

This manuscript describes the development of a module that is part of a learning platform named “NIGMS Sandbox for Cloud-based Learning” https://github.com/NIGMS/NIGMS-Sandbox. The overall genesis of the Sandbox is described in the editorial [NIGMS Sandbox](https://web.uri.edu/riinbre/wp-content/uploads/sites/1497/NIGMS-Sandbox-Manuscript.pdf) at the beginning of this Supplement. This module delivers learning materials on machine learning and decision tree concepts in an interactive format that uses appropriate cloud resources for data access and analyses.

Machine learning (ML) is an important tool in biomedical research and can lead to improvements in diagnosis, treatment, and prevention of diseases. During the COVID pandemic ML was used for predictions at the patient and community levels. Given its ubiquity, it is important that future doctors, researchers and teachers get acquainted with ML and its contributions to research. Our goal is to make it easier for everyone to learn about machine learning. The learning module we present here is based on a small COVID dataset, videos, annotated code and the use of Google Colab or the Google Cloud Platform (GCP). The benefit of these platforms is that students do not have to set up a programming environment on their computer which saves time and is also an important democratization factor. The module focuses on learning the basics of decision trees by applying them to COVID data. It introduces basic terminology used in supervised machine learning and its relevance to research. Our experience with biology students at San Francisco State University suggests that the material increases interest in ML.

## INTRODUCTION

The COVID-19 pandemic made clear how important it is for local governments to collect and analyze epidemiological data for infection control [1–3]. One particular piece of information that was of great importance to governments, healthcare systems and citizens alike was the number of COVID cases at a specific time and location [4–6]. Because of this interest and because of the extensive availability of these types of data, COVID-19 case numbers are a great example to use when teaching data science skills to college students in health-related disciplines. Machine learning has become an important data science technique with many uses in science and medicine [7], but many students in biology and related fields currently do not learn this technology in their regular curriculum. For this reason, we have created a tutorial to teach one important machine learning technique, decision trees, using COVID-19 data as the driving example. We use a small dataset that combines publicly available data from different government websites and research institutions [8–11]. The dataset has been cleaned for the purposes of this tutorial and contains observations from all 58 California counties.

The goal of the tutorial is for students to learn how a decision tree can be used to predict the number of COVID-19 cases per 100,000 people in a county during the summer months of 2020. The variables used as possible predictors for our decision tree are: population size, vaccination rate, unemployment rate and partisan voting rates for different political parties. As the learners work through the tutorial they learn that out of these possible predictors, the unemployment rate is most important for predicting the number of COVID-19 cases. Counties with higher unemployment rates had more COVID-19 cases in 2020, whereas counties with lower unemployment rates had lower numbers of COVID-19 cases. Next, the learners will analyze data from 2021 and discover that in 2021, the best predictor for the number of COVID-19 cases is not unemployment rate, but vaccination rate. Counties with higher vaccination rates had lower numbers of COVID-19 cases while counties with lower vaccination rates had higher numbers of COVID-19 cases. There are a total of four submodules, with code, written text, images and short videos tailored to the needs of a student without coding or machine learning experience.

Decision trees have been used in many research fields for a long time and they are one of the simplest, yet foundational algorithms in machine learning. Decision trees form the basis for random forest models and gradient-boosted tree models, which are some of the most popular machine learning models for supervised learning. Because of its simplicity, the decision tree is a great first algorithm to introduce to college students in order to develop a basic understanding of more advanced techniques within the field of machine learning.

We prepared our tutorial in Python, which is one of the most popular coding languages [12,13] and considered to be a good language to learn for beginners because of its relative simplicity [14]. The tutorial does not require specialized knowledge in any particular biological concept and is designed to be a friendly introduction to the decision tree algorithm for students with no prior coding or machine learning knowledge. We hope that after finishing the tutorial, interested students could use the provided code to analyze other datasets and research questions.

A major barrier for beginners to learn coding is to set up a development environment on their computer. To make sure that many students can use our tutorial, we have provided the code as Google Colab notebooks. Google Colab is a free coding environment included with any free Google account which allows students to get started with coding without needing to set up a development environment on their own computers, and even if they do not own a computer, but instead use a tablet, a Chromebook, their phone or a borrowed computer. We have also created a separate version within the Google Cloud Platform (GCP) for those who already have a GCP account. However, this version of the module requires the user to sign up for a free trial or a paid account. Details on how to access each module will be given in the links in the methods section of this article.

We originally developed the learning module for local biology and chemistry students with limited coding experience and no machine learning experience. We wanted to demystify machine learning concepts and their applications in biology and health-related research. The beginner-friendly learning module aims to get students acquainted with basic terminology used in machine learning with a focus on learning about decision trees. The lessons lean heavily on the practical side, are light on mathematical background and intend to build an intuitive understanding of the algorithm rather than diving into technical and theoretical details. We introduce a few simple best practices, for example, by acquainting students with making comments in the code produced. We teach students how to evaluate their decision tree model and provide them with a positive hands-on experience when they write and run machine learning code for the first time.

The learning module introduces several packages for data wrangling (pandas, numpy), data visualization (matplotlib, seaborn, graphviz) and machine learning (sklearn). All packages are listed in Table 1. These packages are common and used in many other tutorials that are available online, so they provide a good start for students who are interested in learning more.

**Table 1.**
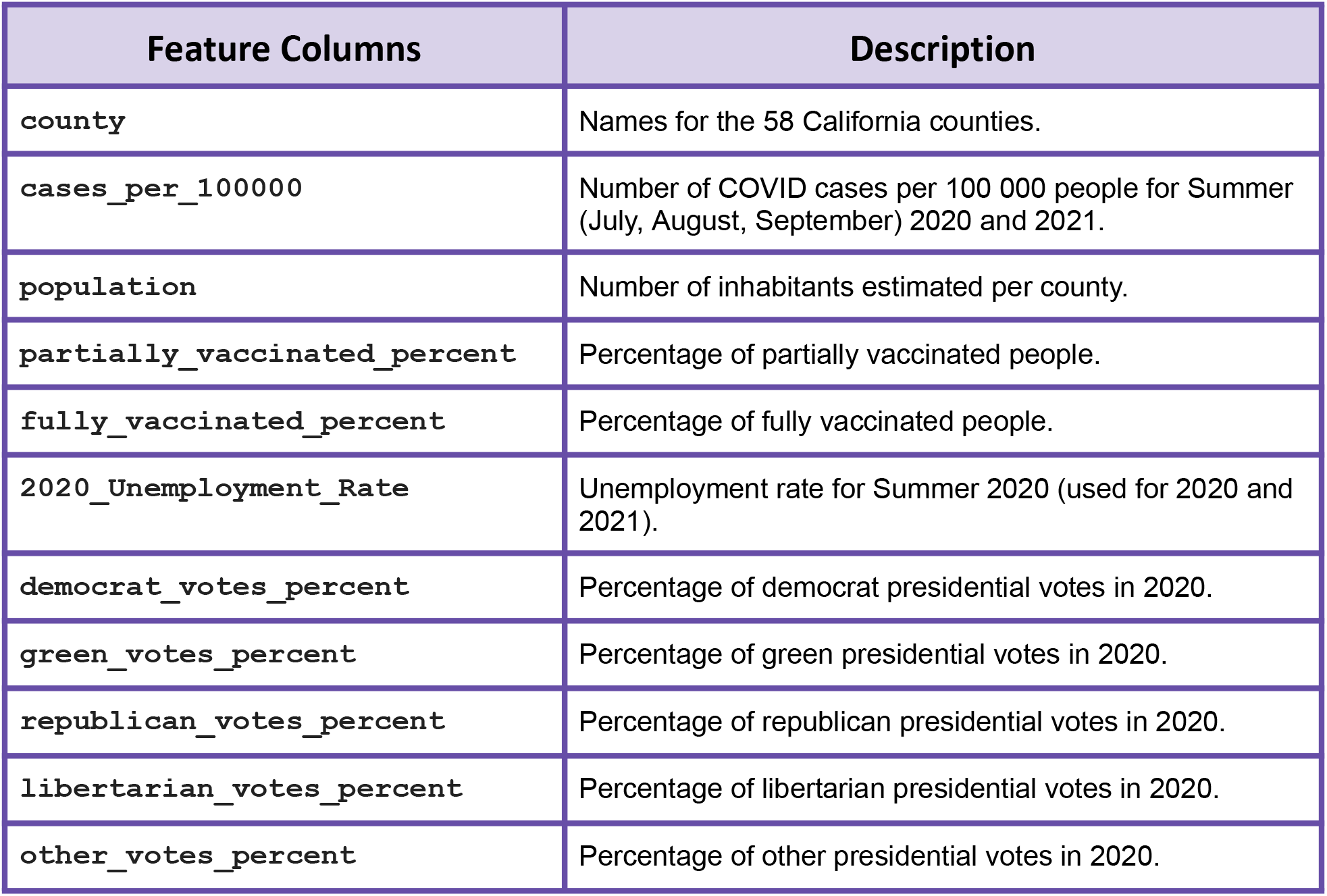
List of variables (features) that are available for each of the 58 California counties.

| Feature Columns | Description |
| --- | --- |
| `county` | Names for the 58 California counties. |
| `cases_per_100000` | Number of COVID cases per 100 000 people for Summer (July, August, September) 2020 and 2021. |
| `population` | Number of inhabitants estimated per county. |
| `partially_vaccinated_percent` | Percentage of partially vaccinated people. |
| `fully_vaccinated_percent` | Percentage of fully vaccinated people. |
| `2020_Unemployment_Rate` | Unemployment rate for Summer 2020 (used for 2020 and 2021). |
| `democrat_votes_percent` | Percentage of democrat presidential votes in 2020. |
| `green_votes_percent` | Percentage of green presidential votes in 2020. |
| `republican_votes_percent` | Percentage of republican presidential votes in 2020. |
| `libertarian_votes_percent` | Percentage of libertarian presidential votes in 2020. |
| `other_votes_percent` | Percentage of other presidential votes in 2020. |

The modules serve to bridge the knowledge gaps between biology students and research practices using machine learning and coding skills. In addition, the submodules are focused on practical use of machine learning and the code can be repurposed for other datasets as well. Students get to learn about the basic workflow used in machine learning from loading the data to evaluating their model performance. We have used the materials presented here for different groups of biology and chemistry students in our classes and summer programs, with encouraging results (see Results section).

The learning module is split into four parts and students can work through one of them or all of them. The first submodule includes videos introducing machine learning and how it is used in research, and then each step of the tutorial is accompanied by a short video explaining what the code is accomplishing. This submodule focuses on the task of predicting the number of COVID-19 cases in counties in 2020 and provides an example of how to create a decision tree, how to visualize it and how to test how well it works. At the end of the module, students will see that the decision tree model works fairly well for 2020, but not for 2021. The second submodule is an optional one where we focus on data drifting and reasons why it is important to retrain our models from time to time. The third submodule provides a notebook for students to practice creating and running the code learned in the first submodule to train a model on the data from 2021. Finally, the fourth submodule is an answer key for the students who are stuck on the third submodule exercise.

## METHODS

The module described in this manuscript is publicly available at https://github.com/MarcMachineLearning/Introduction-to-Machine-Learning (Google Colab version). This link contains the datasets, images used in the notebooks and the collection of tutorial notebooks. To get access to the code the user clicks on the notebook and then clicks the button on top of the notebook that says “*Open in Colab*” within the github repository. This should bring up a new browser tab with a Google Colab notebook open. In that notebook one can then run the code and save a copy to a personal Google Drive.

Those who have access and are familiar with the GCP can follow the instructions on how to access this module here: https://github.com/NIGMS/Introduction-to-Data-Science-for-Biology (GCP version). The GCP version contains some code that only runs correctly within the GCP environment.

The module follows an architecture typical of many data science experiments. Tabular data is available as Comma Separated Values (CSV) files and is read into several analysis pipelines when running the notebooks. After fetching the data, the notebooks demonstrate basic data science functions such as data visualization and exploratory data analysis. Data analysis is executed in notebooks with pre-written code. Each submodule comes with written explanations and videos recorded by the authors.

The learning module was designed with students in mind who have no experience in machine learning and with little or no coding experience. For each step, we explain both the Python code that is used as well as the important machine learning concepts. Short videos, made by the authors, three of whom are graduate students themselves, make the materials more accessible. Here we will describe each of the submodules in more detail.

### Submodule 1: Introduction to Decision Trees

The notebook of the first submodule starts by introducing the data sources used in the tutorial, followed by a video introducing students to the many applications of machine learning in everyday life and in biomedical research (13-minute video). The analysis in this submodule is split into eight steps, and each step is accompanied by a short video (1 to 13 minutes each) explaining the code and some of the concepts used in the notebook. The entire dataset of 58 counties in California, and the number of COVID-19 cases in each of these counties in the summer of 2020 are split into a training dataset of 40 counties and a test dataset of 18 counties. The overall objective of this submodule is to predict the number of COVID-19 cases per 100,000 people in the 18 counties of the test dataset using the variables outlined in Table 1.

After an introduction to machine learning and the project, we outline eight different steps, where code with extensive comments is presented to students to follow along.

1. We show what packages and functions are needed to run the code and how to import them into the notebook to use them.
2. We explain the role of training and testing data in supervised machine learning and import a CSV file with the training data into the notebook.
3. In this step we split the labels or targets (the numbers of COVID-19 cases, which is what we would like to predict) from the features (the data we’ll use to make the predictions).
4. Here we create a decision tree and train it to predict the number of COVID-19 cases per county using the DecisionTreeRegressor function from sklearn.
5. In this step we visualize the resulting decision tree and learn about the different parts of a decision tree.
6. In this step, we introduce the testing data from the 18 counties in California that were not included in the training dataset. We make predictions using the features for these 18 counties and the decision tree we created.
7. Here we provide a graphical and intuitive way to evaluate the performance of the decision tree by loading the actual testing labels (targets) and comparing them with our predictions using bar graphs.
8. Finally in step 8, we use our model, which we trained using data from 2020 to predict the number of COVID-19 cases in the following year, Summer 2021. We see that the model does not perform very well. We also provide a brief discussion as to the reasons why the model performed better when predicting 2020 than with 2021 data within the step 8 video.

At the end of Submodule 1, two CSV files will have been created containing the actual testing labels (targets) and the predictions made for each year by our decision tree. These files will be used in the (Optional) Submodule 2.

### Submodule 2: (Optional) Comparing the model performance for 2020 and 2021

At the end of Submodule 1, the students will have seen that the 2020 model does fairly well when predicting the 2020 COVID-19 numbers, but does not perform as well when predicting COVID-19 numbers for 2021. Submodule 2 revisits this drop in performance of our initial decision tree model and explores some of the reasons for this performance difference. We divided this submodule into five different steps where we explore relationships between the different variables.

1. In step 1, we import packages that focus on visualization and evaluation of our models.
2. Next, we load the CSV files created at the end of Submodule 1, which contain the actual and the predicted values for Summer 2020 and Summer 2021.
3. We then go over the definition and equation of the Root Mean Square Error metric and make a comparison of the predictions made for both years.
4. Here we invite students to suggest some of the reasons why the model performance differs between 2020 and 2021, and we teach them to perform visualizations of these differences through the use of descriptive statistics, histograms, correlation plots and scatterplots with trendlines.
5. Finally in step 5, we introduce the phenomenon of data drift as one of the reasons that machine learning models require retraining from time to time.

### Submodule 3: Independent practice to train a model specifically for 2021

In Submodule 3, we ask students to create a new decision tree using training data from Summer 2021 to improve the performance of the previous model. The submodule contains a video showcasing how this exercise can be performed (10.5 minutes). In this hands-on exercise, we provide a notebook with empty code cells for students to create their own decision tree. Students are encouraged to copy and adapt some of the code from Submodule 1, but they can also try to write the code from scratch if they wish to do so. As a final additional exercise, we ask students to calculate the Root Mean Square Error and create aggregate error histograms, similar to what we have done in Submodule 2.

### Submodule 4: Answer Key for Submodule 3

Submodule 4 offers an answer key for Submodule 3 in case students get stuck or to check the correctness of the answers students came up with.

### Packages and functions used

All code is available as Google Colab Python notebooks. The learning module introduces several packages for data wrangling (pandas, numpy), data visualization (matplotlib, seaborn, graphviz) and machine learning (sklearn). All packages are listed in Table 2. These packages are common and used in many other tutorials that are available online, so they provide a good start for students who are interested in learning more. The same packages are used in the GCP version as well.

**Table 2:** List of packages (or submodules of these packages) and functions or attributes used in the learning module.

| Packages | functions/attributes | Description |
| --- | --- | --- |
| IPython.display | YouTubeVideo() | Displays videos from a YouTube link |
| jupyterquiz | display_quiz() | Displays quiz questions |
| pandas | read_csv()<br>DataFrame()<br>head()<br>drop()<br>concat()<br>plot.barh()<br>reset_index()<br>to_csv()<br>describe()<br>corr()<br>.shape<br>.loc | Read CSV files from a link as a data frame.<br>Organizes data into a 2-dimensional table of rows and columns.<br>Read the first n rows.<br>Remove rows or columns from the data frame.<br>Concatenate pandas objects along a particular axis.<br>Make a horizontal bar plot.<br>Reset the index of the data frame, and use the default one instead.<br>Write object to a comma-separated values (csv) file.<br>Generate descriptive statistics.<br>Compute pairwise correlation of columns, excluding NA/null values.<br>Return a tuple representing the dimensionality of the data frame.<br>Access a group of rows and columns by label(s) or a Boolean array. |
| numpy | arange()<br>triu()<br>ones_like() | Return evenly spaced values within a given interval.<br>Upper triangle of an array.<br>Return an array of ones with the same shape and type as a given array. |
| sklearn.tree | export_graphviz()<br>.fit()<br>.predict() | Export a decision tree in DOT format to be read by graphviz (see below).<br>Trains the algorithm on the training data, after the model is initialized.<br>Given a trained model, it predicts the label (target value) of a new set of data. |
| sklearn.metrics | mean_squared_error() | Calculates the mean square error loss. |
| matplotlib.pyplot | title()<br>hist()<br>legend()<br>show()<br>subplots()<br>set_title()<br>suptitle() | Set a title for one figure plot.<br>Computes and plots a histogram.<br>Place a legend on the axes.<br>Display all open figures.<br>Create a figure and a set of subplots.<br>Set a title for the subplots.<br>Add a centered subtitle to the figure. |
| matplotlib.lines | Line2D() | Draws a line that can have both a solid line style connecting all the vertices, and a marker at each vertex. |
| seaborn | heatmap()<br>light_palette()<br>regplot() | Plot rectangular data as a color-encoded matrix.<br>Make a sequential palette that blends from light to a particular color.<br>Plot data and a linear regression model fit. |
| graphviz | Source() | Imports and reads DOT format files so that the tree can be visualized. |

### Description of the datasets used in the case study

The datasets used in submodules were preprocessed and already separated into training and testing data separately for 2020 and 2021. The datasets consist of COVID case numbers from the California Health and Human Services Agency [8] and COVID vaccination data that were collected by the data desk of the LA Times [11]. We also used unemployment data from the California Employment Development Department [9] and election data from Harvard University [10]. We used unemployment and election data from 2020 for both the 2020 model and for the 2021 model. Voting behavior and unemployment rates were chosen because these data are readily available and we expected them to be correlated with (but not causal to) the number of COVID cases. Machine learning models can lead to accurate predictions, even when they do not lead to an understanding of the underlying mechanisms that lead to the observations of interest. In other words, our model can predict the number of COVID cases for a county based on what the model learned from other counties that have similar characteristics, not because it learns about epidemiology. Students and researchers are encouraged to choose their own features for their own projects and research purposes.

## RESULTS

In 2021 and 2022, we let students at San Francisco State University in the Biology Master’s program and in the summer SCIP program [15] work through the module. We found that our students had limited prior knowledge of machine learning (see Figure 2, panel A), but after doing the tutorial, 71.8% of the students were interested in learning more (the remaining students indicated they were “maybe” interested in learning more, and one student out of 78 indicated they were not interested in learning more). In response to an open-ended question about what they liked about the tutorial some of the answers included: “I liked that the videos were easy to follow and straightforward.”, “I enjoyed getting to change the code myself to try and figure out how things changed in 2021” “Learning to create the decision tree model and learning to create simple visualizations of competing models with the bar plot.”

**Figure1.**
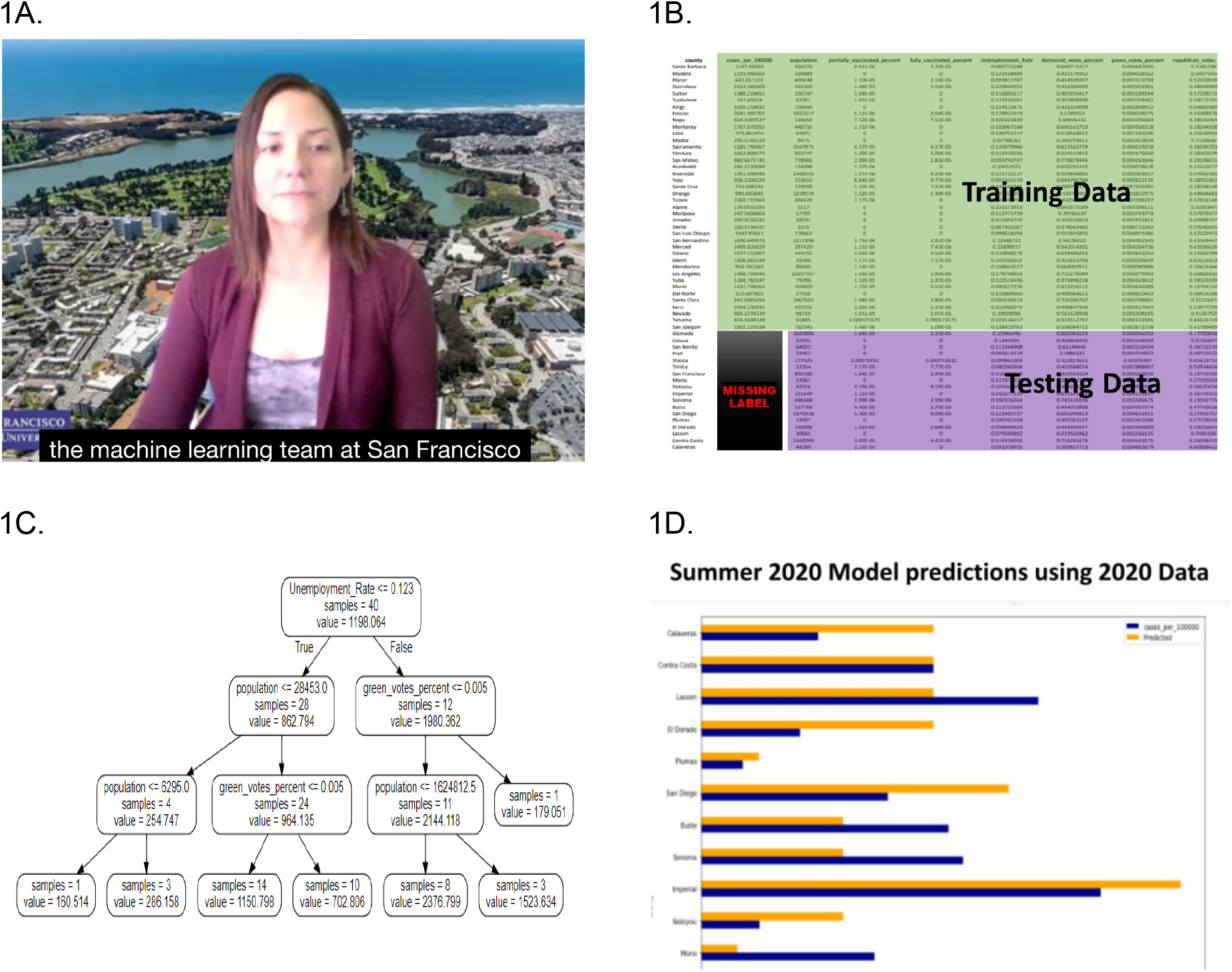
Submodule 1 content **1A**: Still from one of the videos in Submodule 1, where Lorena Benitez Rivera gives a general introduction to machine learning. **1B**: visual explanation of training and testing data. **1C**: The trained decision tree. **1D**: A barplot to compare predicted and actual COVID-19 numbers in 2020.

**Figure 2.**
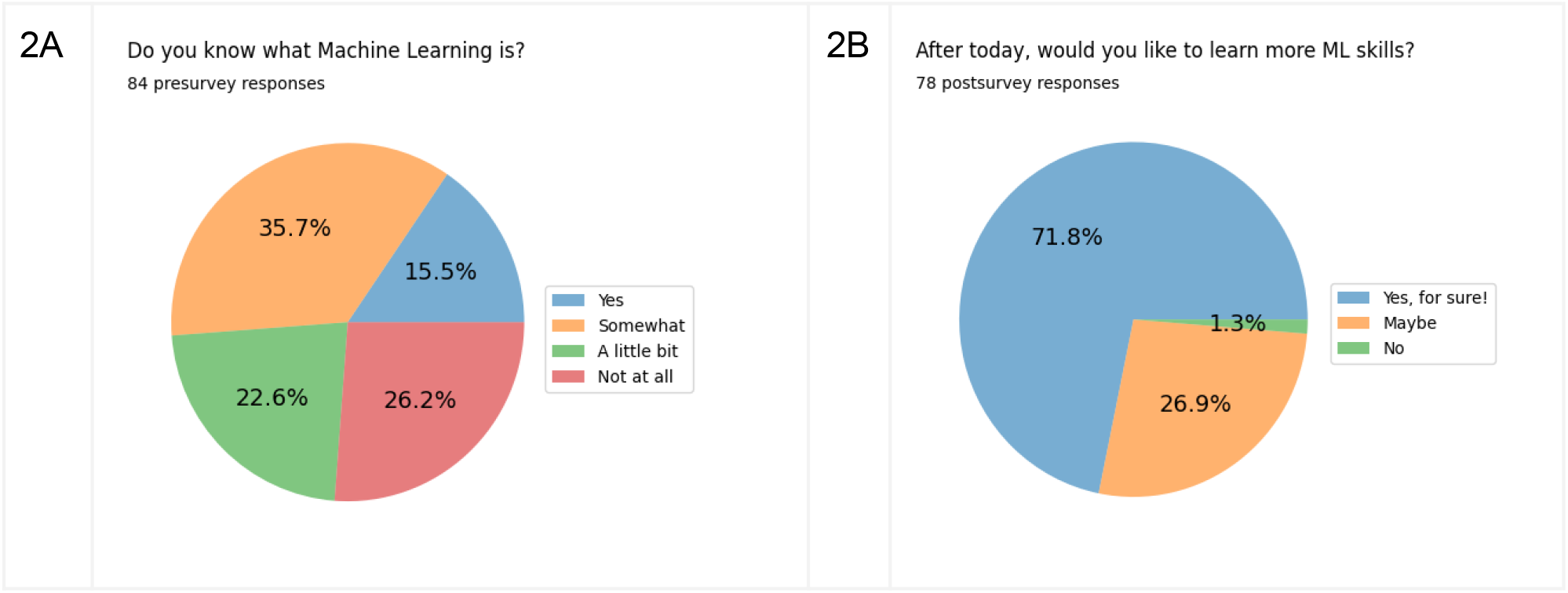
Pre and Post Survey responses. **2A**: Pre-workshop survey responses. **2B**: Post-workshop survey responses. We used the decision tree tutorial in the SCIP summer program and in the Biology Master’s program. Most participants had some coding experience (in R or Python). They had no prior exposure to Machine Learning in their classes or in the Summer program.

## DISCUSSION

Our goal was to create a tutorial that could be used by students with an interest in biology or health, but with no or little coding or machine learning experience. The tutorial is designed such that faculty can have students work through the entire tutorial in a two-hour lab section. Further, we attempted to find a balance between showing students some of the tools they can use for data science and machine learning and helping them learn some of the basic concepts of supervised learning in general and decision trees specifically. We feel strongly that students should learn concepts and techniques at the same time. We hope that the users will realize that both the concepts and the techniques used are simpler than they may have expected beforehand.

Machine learning is a large field and instead of trying to give students an overview of the entire field, we focused here on just one type of supervised learning model, the decision tree. The tutorial should give students a sense of what supervised learning can do. A drawback of this approach is that many other machine learning tools are not discussed in our tutorial. Those who decide to use this tutorial in their lab or classroom may want to clarify that supervised learning is only one part of machine learning and that machine learning is only one sub-discipline of artificial intelligence.

The dataset we focused on in this tutorial is small: 58 observations (one for each county in California), an easy to understand label or target (the number of COVID-19 cases per 100,000) and 9 features (population size, fraction of partially and fully vaccinated, unemployment rate and fraction of votes in the presidential election to different parties). In fact, this dataset is so small it should be considered a toy example. Yet, we picked the topic and the dataset on purpose. Many students in biology are interested in public health and the number of COVID-19 cases was of interest to many in the early years of the pandemic.

In addition to the fact that COVID-19 numbers will be of interest to many biology students, we hope that by using a public health topic for a machine learning tutorial, students will realize that computer science and machine learning can be relevant to topics they care about. To those who are already familiar with machine learning, it may seem obvious that the tools from machine learning can be used in almost any field that is data-heavy, but for biology students, this may not be clear. We were happy to see that a large majority of biology and chemistry students who tried out the tutorial were interested in learning more about machine learning after doing the tutorial (see Figure 2).

Python is a great language for undergraduate students in biology to learn computing skills. Python is one of the most common computer languages at the moment [12,13]. Importantly, it is used often in biology and in computer science departments. The other option for biology students would be R, which also has good data science and machine learning capabilities. However, R is almost never used in computer science departments and is not the best stepping stone for students to get ready to take classes in a computer science department.

The tutorial described here is the result of a collaboration between San Francisco State University, the NIH NIGMS, and a team from Google and Deloitte. This collaboration combined didactic and technical skills of different team members, as well as a variety of backgrounds including biology, computer science and statistics. As a result, the tutorial should be of interest to those who teach college students with a biology background, while it uses approaches and tools that are valued in industry.

The tutorial is available on Github as a set of Google Colab notebooks [17] and also in the Google Cloud Platform [16]. There are several benefits of having students use Google Colab notebooks or the Google Cloud Platform. The most important practical reason is that students can get started more quickly and do not need to install any software on their computers. In fact, students could use a tablet or library computer if they do not own a computer. Because of the availability in the cloud, this material will benefit the research community, especially students and faculty in (Institutional Development Awards) IDeA states and at less research-intensive universities in non-IDeA states.

For this particular tutorial, the barrier to entry would be lowest if students use the Google Colab version, but if they are interested in using the Google Cloud Platform, then this tutorial is available there too. The Google Cloud Platform also allows each student to store their own data in a secure way using Cloud Storage options that prevent data loss and have advanced security options that control data sharing and enable better cross-institutional collaboration. The cloud platform also allows them to have access to many other datasets from official research and/or government institutions to test what they have learned.

Limitations of the approach we took are that at several steps we chose simplicity over completeness. For example, students learn about test and training data, but not validation. Also, students learn about decision trees, but not related methods such as random forests and gradient-boosted trees, and not other common supervised learning methods such as logistic regression, support vector machines, and neural networks. The Google Colab version would currently be easier to access since many students have a Google account.

In conclusion, we hope that the tutorial we have created will be used by many faculty and students to get started with data science and machine learning.

## KEY POINTS

1. Machine learning is an increasingly important field, but it is usually not taught in biology departments.
2. We have created a beginner-friendly introduction to machine learning focused on a simple but relevant COVID-19 dataset from counties in California.
3. The material is made accessible by using cloud-computing platforms which means that all a student needs to participate is a browser window and a Google account.
4. The tutorial includes pre-written code, practice material, an answer key, all accompanied by a series of videos by the makers.
5. Surveys at San Francisco State University showed that after doing the tutorial, biology and chemistry students were interested to learn more about machine learning.

## ACKNOWLEDGMENTS

We are grateful to the many undergraduate and master’s students in the biology and chemistry departments of San Francisco State University who have tested the materials presented here.

## DATA AND CODE AVAILABILITY

Code is available as Google Colab notebooks in this GitHub repository. The code is also available at the NIH GitHub site and data was stored in Google Cloud Storage.

## AUTHOR CONTRIBUTIONS

1. Lucy Moctezuma: Notebook development: coding, testing, graphics, notebook videos, manuscript writing.
2. Megumi Fuse: Conceptualization and supervision of the project.
3. Pleuni Pennings: Conceptualization and supervision of the project. Writing and reviewing the paper.
4. Lorena Benitez: Notebook videos and contribution to manuscript writing.
5. Florentine van Nouhuijs: Notebook videos and contribution to manuscript writing.
6. Faye Orcales: Original data collection and the idea to use California COVID data to teach machine learning.
7. Allen Kim: Notebook development, coding, and testing.
8. Ross Campbell: Notebook development, coding, and testing.

## AUTHOR DESCRIPTIONS

Lucy Moctezuma is a Master’s in Statistics student at CSU East Bay and creates machine learning teaching tools for the Biology and Computer Science departments of San Francisco State University (SFSU).

Megumi Fuse is a biology professor at SFSU and the director of the SEO office which prepares students from underrepresented groups for STEM careers.

Pleuni Pennings is a computational evolutionary biologist at SFSU with a focus on studying drug resistance and on creating ways for biology students to learn quantitative and computing skills. Lorena Benitez is a PINC program alum, GOLD program alum, and Master’s student at SFSU who works on understanding diversity in pathogenic bacteria as well as soil microbiomes on the Galapagos islands.

Florentine van Nouhuijs is a Master’s student and GOLD program alum at SFSU who works on understanding the evolution and transmission of ciprofloxacin resistance in *E. coli*.

Faye Orcales is a PINC program alum (SFSU) and junior research technician at UCSF.

Allen Kim is a Data and AI Engineer at Google. He brings extensive experience in the fields of Machine Learning and Data Science, contributing his expertise to innovative solutions.

Ross Campbell is a bioinformatician at Deloitte with a background in biomarker development from multi-omics experiments. Ross has expertise in optimizing bioinformatics machine learning pipelines to run at scale in cloud environments.

